# Fully resolved assembly of *Cryptosporidium parvum*

**DOI:** 10.1101/2021.07.07.451495

**Authors:** Vipin K. Menon, Pablo C. Okhuysen, Cynthia L. Chappell, Medhat Mahmoud, Medhat Mahmoud, Qingchang Meng, Harsha Doddapaneni, Vanesa Vee, Yi Han, Sejal Salvi, Sravya Bhamidipati, Kavya Kottapalli, George Weissenberger, Hua Shen, Matthew C. Ross, Kristi L. Hoffman, Sara Javornik Cregeen, Donna M. Muzny, Ginger A. Metcalf, Richard A. Gibbs, Joseph F. Petrosino, Fritz J. Sedlazeck

## Abstract

**Background:** *Cryptosporidium parvum* is an apicomplexan parasite commonly found across many host species with a global infection prevalence in human populations of 7.6%. As such, it is important to understand the diversity and genomic makeup of this prevalent parasite to fight established infections and prohibit further transmission. The basis of every genomic study is a high quality reference genome that has continuity and completeness, thus enabling comprehensive comparative studies.

**Findings:** Here, we provide a highly accurate and complete reference genome of *Cryptosporidium parvum*. The assembly is based on Oxford Nanopore reads and was improved using Illumina reads for error correction. We also outline how to evaluate and choose from different assembly methods based on two main approaches that can be applied to other *Cryptosporidium* species. The assembly encompasses 8 chromosomes and includes 13 telomeres that were resolved. Overall, the assembly shows a high completion rate with 98.4% single copy BUSCO genes.

**Conclusions:** This high quality reference genome of a zoonotic IIaA17G2R1 *C. parvum* subtype isolate provides the basis for subsequent comparative genomic studies across the *Cryptosporidium* clade. This will enable improved understanding of diversity, functional and association studies.

## Introduction

*Cryptosporidium* is an apicomplexan parasite of public health and veterinary significance with a recent analysis reporting a global infection prevalence of 7.6% [1]. Historically, limited government and private funding was available to study the epidemiology and molecular dynamics of the organism, but this has recently shifted [2].

*Cryptosporidium* spp. have been found in 155 species of mammals, including primates [3,4]. Among humans, twenty species of *Cryptosporidium* spp. have been identified [5]. Although the parasite can be transmitted in a variety of ways, the most common method is *via* drinking and recreational waters. In the United States, *Cryptosporidium* is the most common cause of waterborne disease in humans [6]. Studies have shown that *Cryptosporidium* is responsible for a large proportion of all cases of moderate-to-severe diarrhea in children under the age of two [7,8]. There is currently no vaccine available, and the only approved drug for the treatment of *Cryptosporidium*-related diarrhea is nitazoxanide (NTZ), which has limited activity in immunocompromised patients.

Previously, the inability to complete the life cycle of *Cryptosporidium in vitro* hampered progress in understanding pathogenesis and exploring new treatment modalities. Recent advances using human organoids support the full parasite life cycle, recapitulate *in vivo* physiology of host tissues [9][10–12], and provide a way to study the molecular mechanisms and pathways used by *Cryptosporidium* during infection. However, to facilitate genomic or association studies, a high quality reference genome is needed.

*C. parvum* was included in early genome-sequencing projects due to its public health importance and high global prevalence. The first reported complete genome assembly for *C. parvum* Iowa II became available in 2004 [13], generated by random shotgun sequencing approach, resulting in roughly 13x genome coverage totaling 9.1 Mb of DNA sequence across all eight chromosomes. This reference sequence had a reduced coverage across the genome, with multiple gaps and was not adequate to represent the full breadth of genes present, which could result in misleading interpretations of the isolates being studied. In addition, online repositories such as GenBank, CryptoDB and the Wellcome Trust Sanger Institute FTP servers provide a range of unassembled, unprocessed raw read sequences.

Long-read sequencing technology has advanced to enable read lengths of 15 - 20 Kb (PacBio) and 2 - 3 Mb (Oxford Nanopore (ONT)) with low error rates and is frequently utilized to improve reference genome assembly [5,14–19], thus, enabling long continuous assemblies without gaps even across highly repetitive regions [20]. While long-read technologies enable an improved assembly, it is difficult to evaluate which *de novo* assembly best represents the sample. Currently, the simplest way to rank *de novo* assemblies is by length [20] (N50) or BUSCO [21] comparison. However, this is not a guarantee that chromosomes are well represented or correctly arranged. Furthermore, the variety of *de novo* assembly methods (Canu [22], Flye [23], Shasta [24], Falcon [25], etc.) makes it harder to choose the best representation.

In the current study, we have generated a reference genome for *C. parvum* by using long-read sequencing on the ONT PromethION supplemented with short-read data generated on NovaSeq 6000 for error correction (see **Figure 1**). This resulted in a complete reference including all chromosomes and thus represents a gap-less representation of this important pathogen. Furthermore, it includes 13 of 16 telomeric sequences. The assembly is available at PRJNA744539 (GCA_019844115.1). In addition to the novel assembly, we lay out our QC process and assessment of the assembly to optimize not only for length but also to assess the overall structure of the draft assemblies. Following this comparison schema, it is easy to choose the most optimal representation. In addition, this schema is applicable for other species as well, from single haploid to more complex organisms like plants or humans.

**Figure 1:**
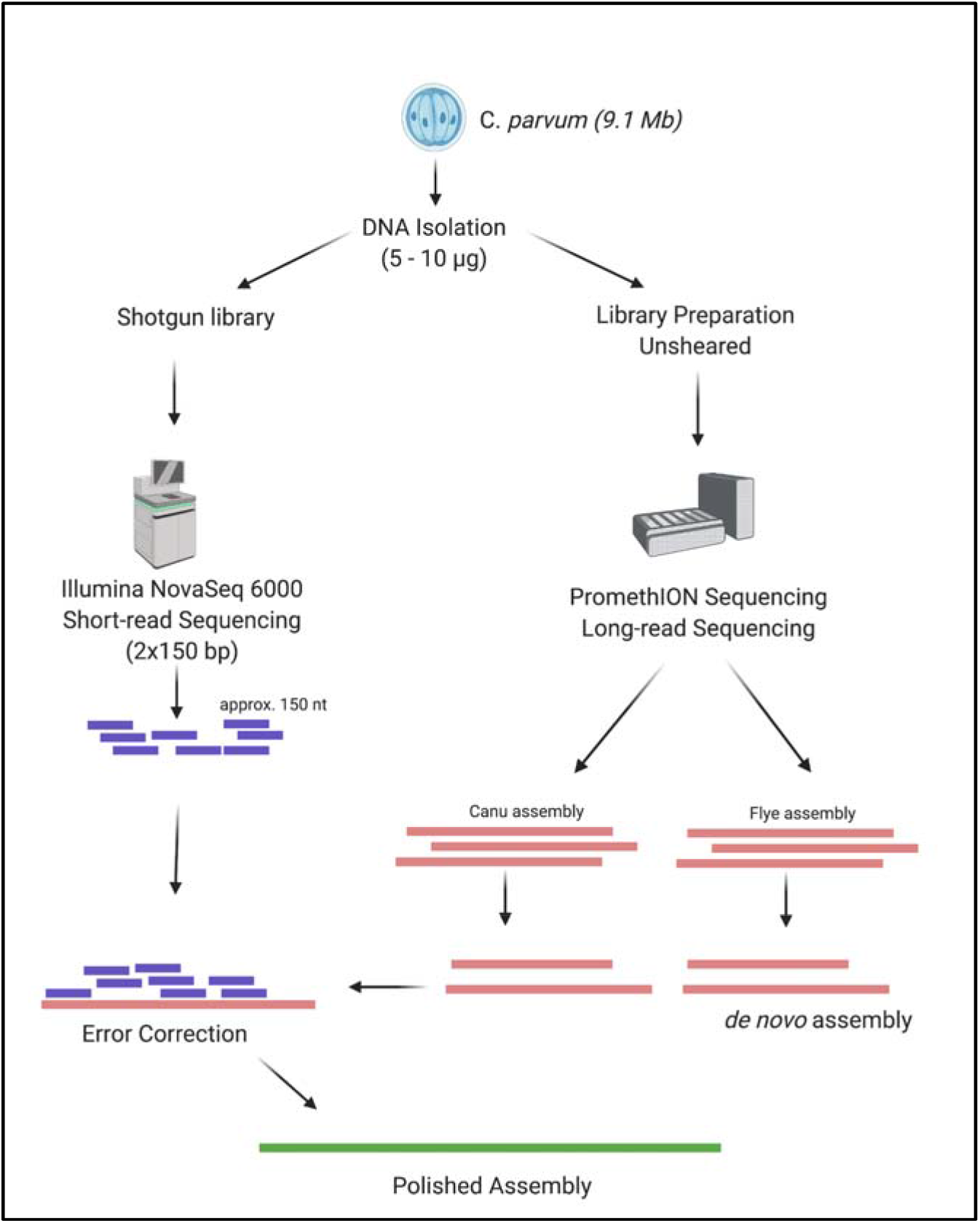
Workflow for the generation of Cryptosporidium parvum assembly.

## Results

We sequenced the *C. parvum* genome with Oxford Nanopore long-reads (see methods) and obtained a total of ~480Mbp of sequence (**Figure 1**). This is equivalent to 53x coverage for this genome (~9Mbp genome size). **Figure 2** shows overall statistics on read length and coverage. The N50 read length is 15.3 kbp with 10x coverage of reads with ≥30kbp length. Our longest read detected was 808 kbp. In addition, we sequenced the genome using the Illumina NovaSeq 6000 to produce 352x coverage of 150 bp paired end reads.

**Figure 2:**
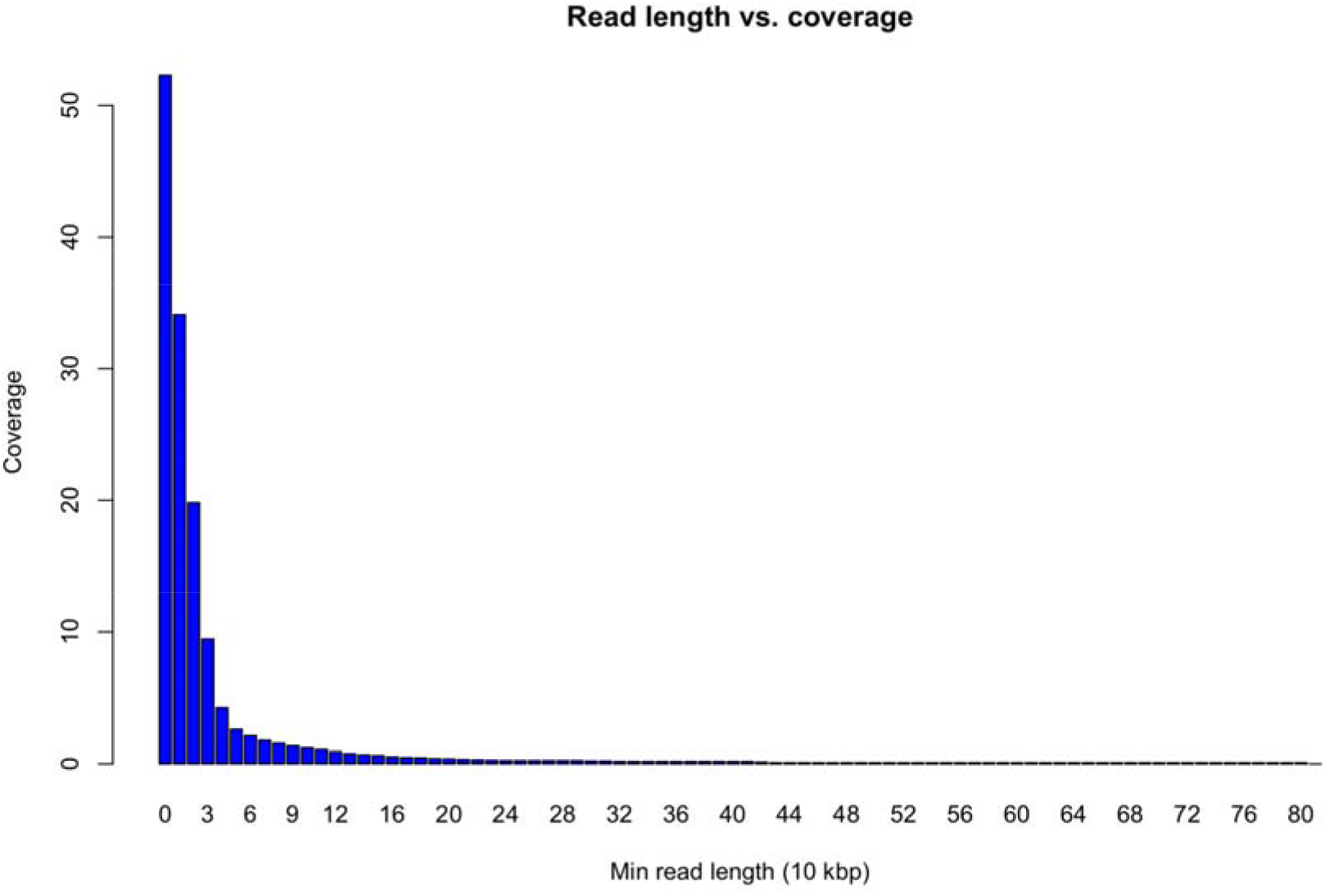
Read length distribution and cumulative coverage over the Oxford Nanopore. Sequencing. We obtained a total of 53x coverage with long-reads and even 10x coverage with reads larger than 30kbp (x axis). The longest read measured was 808kbp.

Using these short-reads we ran a genome estimation using GenomeScope [26] to obtain a genome size estimate using a polyploidy of 1. Doing so resulted in an estimate of 9.9Mbp with an 89.24% model fit (see **Supplementary Figure 1**). Inspection of the resulting data (Figure 2) highlights that this is a potential overestimation of the genome size itself and thus fits in the realm of the previously reported reference assembly in CryptoDB (GCA_015245375) of ~9.1Mbp.

### Assembly and comparison of *Cryptosporidium* Assembly

The initial assembly was carried out with only the ONT reads using Canu [22] (see methods) and resulted in 25 contigs with 8 contigs representing all chromosomes. We obtained a total genome length of 9.19Mbp across 8 assembled contigs with an average N50 size of 1.11Mbp (Table 1). The largest contig was 1.4 Mbp. Our assembly shows a NG50 similar to that of the assembly published in 2004 (see **Figure3 A**).

**Table1:** Overall assembly statistics and comparison using Quast between the current assembly and the previously established assembly.

|  | **GCA_000165345.1** | **GCA_019844115.1**<br>**(newly established)** |
| --- | --- | --- |
| Total sequence length | 9,102,324 | 9,197,619 |
| Total ungapped length | 9,087,655 | 9,197,619 |
| Unresolved sequences | 14,669 | 0 |
| N50 | 1,104,417 | 1,108,772 |
| N90 | 985,969 | 993,129 |
| L50 | 4 | 4 |
| Total number of chromosomes | 8 | 8 |

**Figure3:**
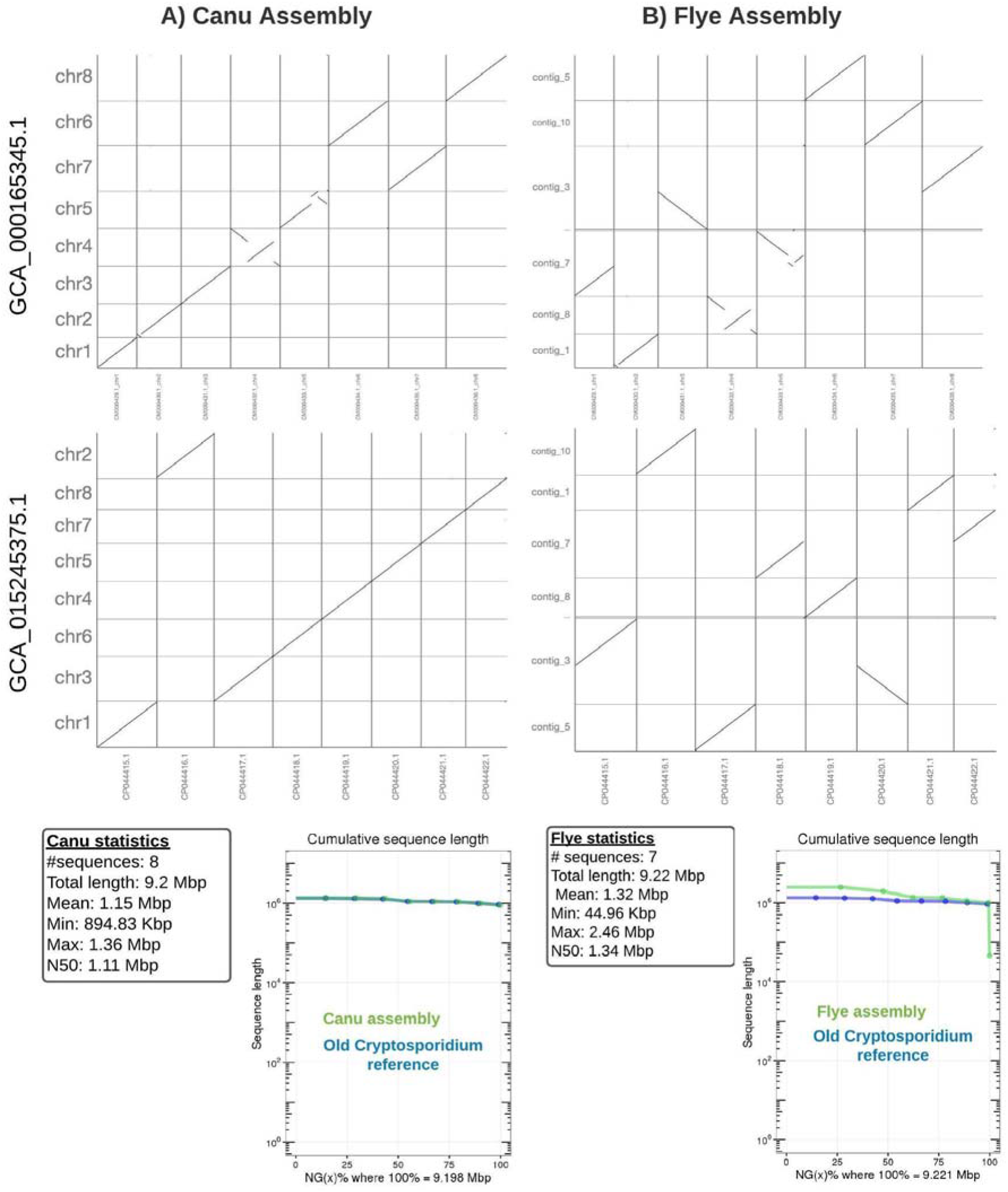
Assembly comparisons. A) The Canu assembly shows a high concordance with the previously published C. parvum assembly (GCA_015245375.1) [27] (dotplots) and agreements in length (bottom). Nevertheless clear assembly differences are visual when comparing it to GCA_000165345.1 [13] B) The Flye assembly versus the C. parvum assembly (GCA_015245375.1) shows large disagreements. Contig 3 is merged between two different Cryptosporidium chromosomes, and one chromosome is missing. Also, the length comparison (bottom) shows discrepancies in the beginning highlighting a very short contig in the end (green track). Interestingly GCA_000165345.1 shows structural differences over both assemblies likely indicating errors in the previous reference.

We also generated an assembly with Flye assembler[23] (see methods), which led to a total of 7 contigs. However, one contig was only 62,160 bp long (see **Figure3 B**). Despite this early warning sign, we compared the two assemblies to identify which one best represented the *C. parvum* genome using genome alignments and remapping of short reads.

To validate our findings, we first aligned the Canu and Flye assemblies to the previously published *C. parvum genome* reference [3] using nucmer [28](v3.23). The nucmer alignments were filtered by “-l 100 -c 500 -maxmatch” for all assemblies following the suggestions from Assemblytics [29], which was used to study the alignment results that were generated (Figure 3).

The dot plot from a MUMmer alignment analysis indicates that the GCA_015245375.1 [27] and Canu genome assemblies are largely collinear (**Figure 3A**). All chromosomes show co-linearity to the previously established assembly for *C. parvum*. Upon closer inspection small segments that aligned to other chromosomes were shown to be telomeric sequences. Thus, these segments did not indicate inaccurate alignments *per se*, but highlighted their repetitive nature (see below for details on telomere reconstruction). However, when assessing the dot plot generated for the Flye assembled genome(**Figure 3B**), we observed larger disagreements compared to GCA_015245375.1. As previously mentioned, one contig from the Flye assembly was small (62 kbp) and judged to be an artifact. More problematic, however, was the merger of two *Cryptosporidium* chromosomes into contig3 (**Figure 3B**, second to last row in dotplot). A fusion of two chromosomes from *Cryptosporidium* was also observed on contig_7. Overall, these analyses show that while we initially missed one contig (7 instead of the expected 8), which was too small (~62kbp) to represent a chromosome. Thus, the missing two chromosomes were merged with other chromosomes within two contigs from Flye. When comparing both of our assemblies (Canu and Flye) to the previously established GCA_000165345.1, we saw large structural disagreements on both assembly comparisons (**Figure 3A/B**). The differences between GCA_000165345.1 and our *de novo* assemblies are most likely due to structural faults in GCA_000165345.1.

We further carried out a remapping experiment to identify structural disagreements between the Illumina data (short-read) and the long-read assemblies. We mapped the reads and found structural variants (SVs) based on discordant paired end reads (see methods)[30]. We identified a total of 10 potential SVs over the remapping based on the Flye assembly. The majority of events were insertions (4) followed by duplications (3) and breakend (BND) (2). However, on closer inspection only two SVs (the two BND) showed a misassembly with a homozygous alternative genotype. All other eight SVs showed a minor allele frequency and are likely consequences of mapping artifacts or heterogeneity of the sequenced population. Next, we assessed the Canu assembly, which showed nine SVs in total. All of the identified SVs showed a low read support, indicating a low probability of being correctly identified and likely originating from mapping artifacts as the material originates from a pure oocyst (see methods). This assessment demonstrated that the Canu assembly is the better representation of *C. parvum* compared to Flye assembly for this study.

### Establishing *Cryptosporidium* Assembly

The quality of the Canu generated draft assembly was further improved by two rounds of assembly polishing employing the short-reads (see methods). After the first round of polishing, the number of corrections were reduced to ~20 along the entire genome. Eight largest contigs available in the final polished assemblies are aligned (see methods) to the previously published *C. parvum* reference GCA_015245375.1 [13]. The alignment analysis further confirmed that the eight contigs represent the previously published chromosomes, while the other contigs appear to be repeats at the start or end of the contigs. Our assembled eight chromosomes complete 14,669 bp of unresolved sequences (i.e. N). Our assembly also showed a GC content (30.11%) similar to the previous version (30.18%), again attesting to the overall quality.

To further assess the completeness of our assembly, we used Busco [21] with the coccidia_odb10 linkage set (see methods). This analysis confirmed the high quality of our assembly, showing 494 (98.4%) complete re-identified genes from a total of 502. All 494 genes had single copies, indicating that the new assembly is error-free. In addition to these single-copy genes, three genes were fragmented, and five genes were missing from the Busco run.

A further comparison to the previous reference genome (GCA_015245375.1) [13] revealed a high consistency with only four structural variants (one insertion, one deletion, one tandem expansion and one tandem contraction) between the two assemblies. This comparison was done based on the genomic alignment and using Assemblytics [29].

Lastly we used the Illumina data set to identify SNV with respect to the new assembly (GCA_019844115.1). **Supplementary Figure 2** shows the allele frequency of the passing SNV (see methods) and indicates that there are no major differences to be observed and also highlights the purity of the utilized material for the assembly process.

### Telomere identification

Telomeric ends present on either end of each chromosome were identified in the Canu genome assembly (see methods). To search for telomeres, we identified matching sequences of “TTTAGG” repeats [31] in our assemblies (see methods). Telomeric areas were defined as those with at least 100 repeated sequence matches within a region near the start and end of the contigs. Given these conservative thresholds, we identified a total of 13 telomeric regions. For the majority of chromosomes (2,3,4,5 and 6) telomeric regions were identified at both ends of the chromosomes, thus fully representing the chromosomes from telomere to telomere, including the centromere. telomeres were only observed at the beginning of chromosomes 7, 1 and at the end of chromosome 8. We further-crossed checked the other contigs that were previously filtered out. These highlighted telomeric sequences, but couldn’t be placed automatically to the other chromosomes (i.e, chromosomes 1, 7 or 8). Overall, the identification of the telomeric sequences on the vast majority of the contigs highlights the overall high quality and continuity of our newly established *C. parvum* genome. The final assembled genome has been deposited at GenBank (accession GCA_019844115.1).

### Assessment of subtyping loci

*Cryptosporidium* spp. Are usually typed and characterized widely by using a small set of genetic markers including *gp60*, COWP, HSP70 and 18S [32]. Most of the genetic marker data available in GenBank were generated from short-read amplification and sequencing by Sanger, thus providing an improved resolution, but still contain errors arising from manual curation.

The *gp60* sequence from the current assembly was aligned with reference sequences retrieved from GenBank. Reference sequences selected for alignment consisted of multiple IIa (*C.parvum*) subtypes, including a IIaA17G2R1 reference (MK165989) corresponding to the sequenced C. parvum isolate in our study. ClustalW alignment was carried out using BioEdit V7.2.5 With no gaps or large mismatches. The assembled genome has 100% identity with the reference genome IIaA17G2R1, and the genetic markers were observed (see **Supplementary Figure 3**).

## Conclusion

The current work highlights how next-generation sequencing, including third-generation long-read sequencing, can be used to generate a high-quality genome assembly complete with centromeric regions and numerous telomeres. The genome assembly generated provides a gapless reference compared to the previously published GCA_000165345.1 [13] and extends into some telomeric regions over GCA_015245375.1 [27]. Telomeric regions added to those from GCA_000165345.1, which is a hybrid assembly based on two different subtypes of *Cryptosporidium spp*. (IIaA17G2R1 and IIaA15G2R1), which might impact further comparison or association studies. In contrast, our study was able to boost the fidelity and robustness of the assembly by focusing on one subtype only, IIaA17G2R1 resulting in a better telomere to telomere assembly representation (GCA_019844115.1). Studies of *Cryptosporidium spp. are* based on genetic markers previously identified for some regions of chromosome 6, and are not able to provide a better understanding of the genetic variation and recombination occurring within the species. Thus, establishing stronger marker genes and perhaps enabling improved recovery of *Cryptosporidium*-specific sequencing reads by mapping to a high-resolution reference genome will enable better understanding of *Cryptosporidium* transmission.

A commonly used approach for *C. parvum* subtyping is based on tandem repeat analysis of *gp60*, a highly polymorphic gene that encodes for an immunodominant glycoprotein (15/40 kDa) located on the surface of sporozoites and merozoites of many *Cryptosporidium* species[33]. The current study was done using an isolate propagated in calves by Bunch Grass Farms (Deary, ID). The vendor originally propagated *C. parvum* IOWA II belonging to subtype IIaA15G2R1 based on *gp60* sequencing. This strain has now been replaced with a closely related local isolate belonging to the IIaA17G2R1 subtype. In our work, this isolate is referred to as *C. parvum* (GCA_019844115.1). It is unclear if the IIaA17G2R1 evolved from IOWA II, possibly from recombination with another local isolate, or if it represents a distinct isolate on its own. To our knowledge the assembly done here represents the first IIaA17G2R1 subtype isolate for which long read sequencing has been performed. *C. parvum* isolates belonging to the IIaA17G2R1 subtype have been identified in farms in various regions of the world[34–36], was the second most common genotype identified in human cases in a recent study done in Canada[37] and is responsible for causing foodborne outbreaks in the US[38,39]

Published studies have shown the presence of contingency genes in *Cryptosporidium* spp., which are responsible for surmounting challenges from the host and are subject to spontaneous mutation rates [40–42]. The majority of these genes are located in the telomere regions of the chromosomes, which are prime sites that evolve and mediate host-parasite interactions [31,43]. In the current assembly, we were able to resolve 13 of the estimated 16 telomeres. The capacity to resolve telomeres and subtelomeres across chromosomes in *Cryptosporidium* spp. will lead to a better understanding of the organism’s adaptation to a variety of environmental and host settings.

We utilized two *de novo* assembly approaches here to obtain a better representation for *Cryptosporidium* spp. and demonstrated two methods for validating these two assemblies. First, we compared the assemblies from Flye and Canu to pre-existing assemblies from *Cryptosporidium* spp. from different subtypes and were able to identify certain structural differences. Further, the detection of structural variations (SVs) proved very helpful in deciding which assembly best represents the species at hand[20]. This was only possible by having orthogonal sequenced Illumina reads. Other studies might choose a different strategy such as utilizing HiC directly, which would also enable a better scaffolding [44]. For *Cryptosporidium* spp this was not necessary as the genome is of relatively small size (~9Mbp) and encompasses eight chromosomes. The analysis of Busco is also a very important indication of quality (i.e., completeness and redundancy) but didn’t indicate incorrect rearrangements identified with the Flye assembly. These types of mis assemblies can be readily identified only by comparing closely related reference genomes and/or orthologous data sets (e.g., Illumina short reads)..

The final *Cryptosporidium* spp. assembly will be a helpful resource to advance the study this important pathogen, further investigate its complexity during growth and development *in vitro*, and serve as a reference for the study of genetic diversity among different isolates. Furthermore, we hope it also facilitates translational research that focuses on characterizing virulence, pathogenicity, and host specificity. In this way, new targets may be found leading to vaccines or effective antiparasitic agents to treat this important pathogen.

## Methods

### DNA extraction

*Cryptosporidium parvum* oocysts were obtained from Bunchgrass Farm in Deary, ID (Lot #22-20, shed date, 10/2/20) and are propagated from IOWA-1 subtype IIaA15G2R1, which was recently replaced by a local isolate subtype IIaA17G2R1[45]. Purified oocysts (10^8^) were washed in PBS and treated with diluted bleach for 10 minutes on ice to allow for sporozoite excystation. Parasites were pelleted, washed in PBS, and DNA was extracted using Ultrapure™phenol:chloroform:isoamyl alcohol (Thermo Scientific) followed by ethanol precipitation. Glycoblue™ co-precipitant (Thermo Scientific) was used to facilitate visualization of DNA during extraction and purification steps.

### ONT Library preparation & sequencing

NEBNext FFPE DNA Repair Mix was used to repair 620ng of genomic DNA, which was then followed by end-repair and dA-tailing with NEBNext Ultra II reagents. The dA-tailed insert molecules were further ligated with an Oxford Nanopore adaptor via ligation kit SQK-LSK110. Purification of the library was carried out with AMPure XP beads (Beckman, Cat# A63880), the final library of 281ng was loaded to one PromethION 24 flow cell (FLO-PRO002) and the sequencing data was collected for 24 hours.

### Illumina Library preparation & sequencing

DNA (100 ng) was sheared into fragments of approximately 300-400 bp in a Covaris E210 system (96 well format, Covaris, Inc. Woburn, MA) followed by purification of the fragmented DNA using AMPure XP beads. DNA end repair, 3’-adenylation, ligation to Illumina multiplexing dual-index adaptors, and ligation-mediated PCR (LM-PCR) were all completed using automated processes. The KAPA HiFi polymerase (KAPA Biosystems Inc.) was used for PCR amplification (10 cycles), which is known to amplify high GC and low AT rich regions at greater efficiency. A fragment analyzer (Advanced Analytical Technologies, Inc) electrophoresis system was used for library quantification and size estimation. The libraries were 630 bp (including adaptor and barcode), on average. The library was pooled with other internal samples, with adjustment carried out to yield 3 Gbp of data on a NovaSeq 6000 S4 flow cell.

### Genome size estimation

We used Jellyfish (version 2.3.0) to generate a k-mer based histogram of our raw reads in order to estimate the genome size based on our short read data. To obtain this we ran Jellyfish[46,47] with “ jellyfish count -C -m 21 -s 1000000000 -t 10” and subsequently the “histo” module with default parameters. The obtained histogram was loaded into GenomeScope[46] given the appropriate parameter (k-mer size of 21) and haploid genome. GenomeScope provided the overall statistics across the short reads.

### Assembly evaluation

We aligned the assembly of Canu (version 2.0)[22] and Flye (version 2.8.1-b1676) [23] with the two Cryptosporidium assemblies GCA_000165345.1 and GCA_015245375.1 using nucmer (version 3.1) -maxmatch -l 100 -c 500 [28]. Next, the delta files were evaluated with Assemblytics[29] (version 1.2.1) (assemblytics.com) using the dotplot function. In addition, we mapped the short Illumina reads using bwa mem[48] (0.7.17-r1188) with default parameters to our new assembly. Subsequently, we identified structural variants using Manta[49] (v1.6.0) and assessed the VCF file manually. Manta identifies SV based on abnormally spaced or orientated paired end illumina reads here with respect to our new assembly. We further assessed the Illumina data by identifying SNV using iVar[50] (version 1.3.1) with default parameters (cite). We summarized the allele frequencies across the reads using a custom bash script for PASS variants only

### Assembly and polishing

We utilized Canu [21] (v2.0) for the assembly, which was based only on Nanopore pass data and a genome size estimate of 9Mbp. On the Nanopore pass reads, we also ran the assembly using Flye [22] (version 2.8.1-b1676) with the default parameters. Subsequently, we aligned the short reads using bwa-mem (version 0.7.17-r1188) with -M -t 10 parameters. Samtools[51] (v1.9) was used to compress and sort the alignments. The so generated alignment was used by Pilon [52] (v 1.24) with the parameters “--fix bases “by correcting one chromosome after another of the raw assembly. This process was repeated two times achieving a high concordance of the reads and the long-read assembly at the 2nd polishing step.

### BUSCO assessment

We ran BUSCO [21] (v5.2.2) to assess the completeness of our assembly using the parameter “busco-m geno-l coccidia_odb10 -i”, coccidia_odb10 (Creation date: 2020-08-05, number of genomes: 20, number of BUSCOs: 502). The summary statistics generated by Busco are presented under results.

### Telomere Identification

We used the sequence “TTTAGGTTTAGGTTTAGG” to identify telomeric sequences at the start and end of every contig from our assembly. To do so we used Bowtie[53] (version 1.2.3) to align the telomeric sequence back to the assembly with -a parameter. Subsequently we counted the matches across regions using a custom script. In short, we used 10kbp windows to count the number of reported hits, align the genome and compare the locations with the expected start/end locations. The identified regions were filtered for at least 100 hits to guarantee a robust match. This way, we counted the number of times each chromosome was listed.

### Regional comparison

Genetic marker gp60 was used to subtype the assembled genomeagainst available GenBank reference genomes for *C.parvum*. Representative reference genomes for *C. parvum* were downloaded from GenBank and were aligned using ClustalW [54](BioEdit V7.2.5) against the current assembly. Further analysis of the *gp60* gene sequence for tandem repeats to determine subtype designation was done following the methods of Alves et. al. [55]

## Additional Files

Supplemental Figure 1. Genomescope estimation of genome size

Supplemental Figure 2. ClustalW alignment of the *gp60* coding sequence with the assembly.

## Competing Interests

The corresponding author of the paper has presented at both ONT and PacBio sponsored conferences.

## Funding

This work was supported by the National Institute of Allergy and Infectious Diseases (Grant#1U19AI144297).

## Authors’ Contributions

F.J.S and V.K.M : Conceptualization, Analysis and Writing-Original Draft Preparation

C.C and G.A.M : Conceptualization and Writing-Review & Editing

P.C.O : Conceptualization, Resources and Writing-Review & Editing

H.D.; Q.M. and D.M.M. : Conceptualization, Writing-Review & Editing

S.S.; S.B.; K.K.; G. W.; H.S.; V.V.; Y.H. : Methodology, Investigation

M.C.R.; K.L.H.; S.J.C. : Conceptualization

M.M; M.M..: Analysis

R.A.G.; J.F.P. : Conceptualization, Funding Acquisition

## Notes

### Competing Interest Statement

The corresponding author of the paper, Fritz J. Sedlazeck has presented at ONT and PacBio sponsored conferences.

### Summary of Updates

In response to the reviewers, changes have been incorporated to different sections of the paper (Methods and Discussion).

