## Supplemental Information for "Fully resolved assembly of *Cryptosporidium parvum*"

### Supplement

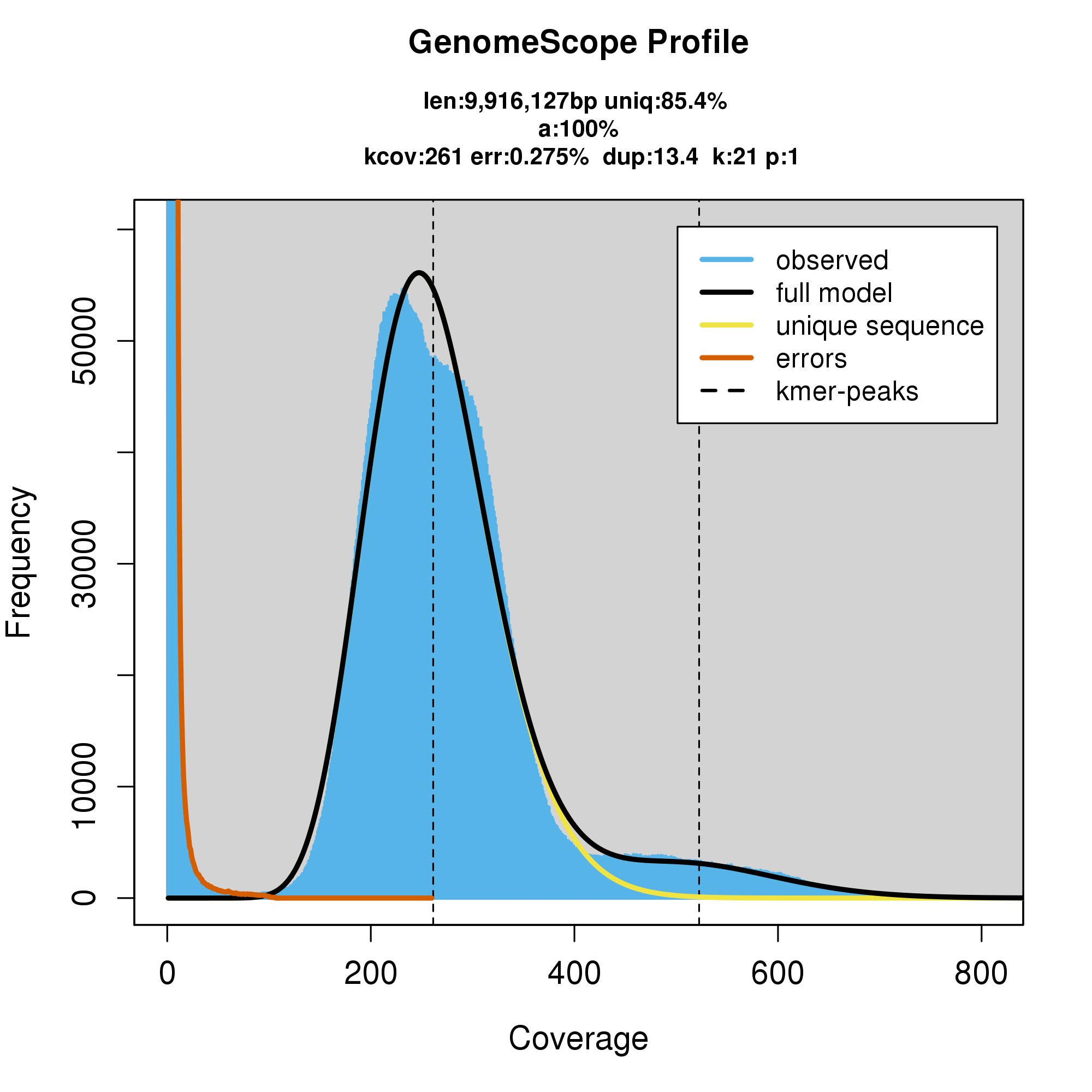

**Supplementary Figure 1:** Genomescope estimation of genome size. The model predicted a genome size of 9.9 Mbp, but as one can see the model fit is only 85.4% not well representing the tip of the variant (black) peak. Thus likely the estimation of 9.9Mbp is a maximum size.

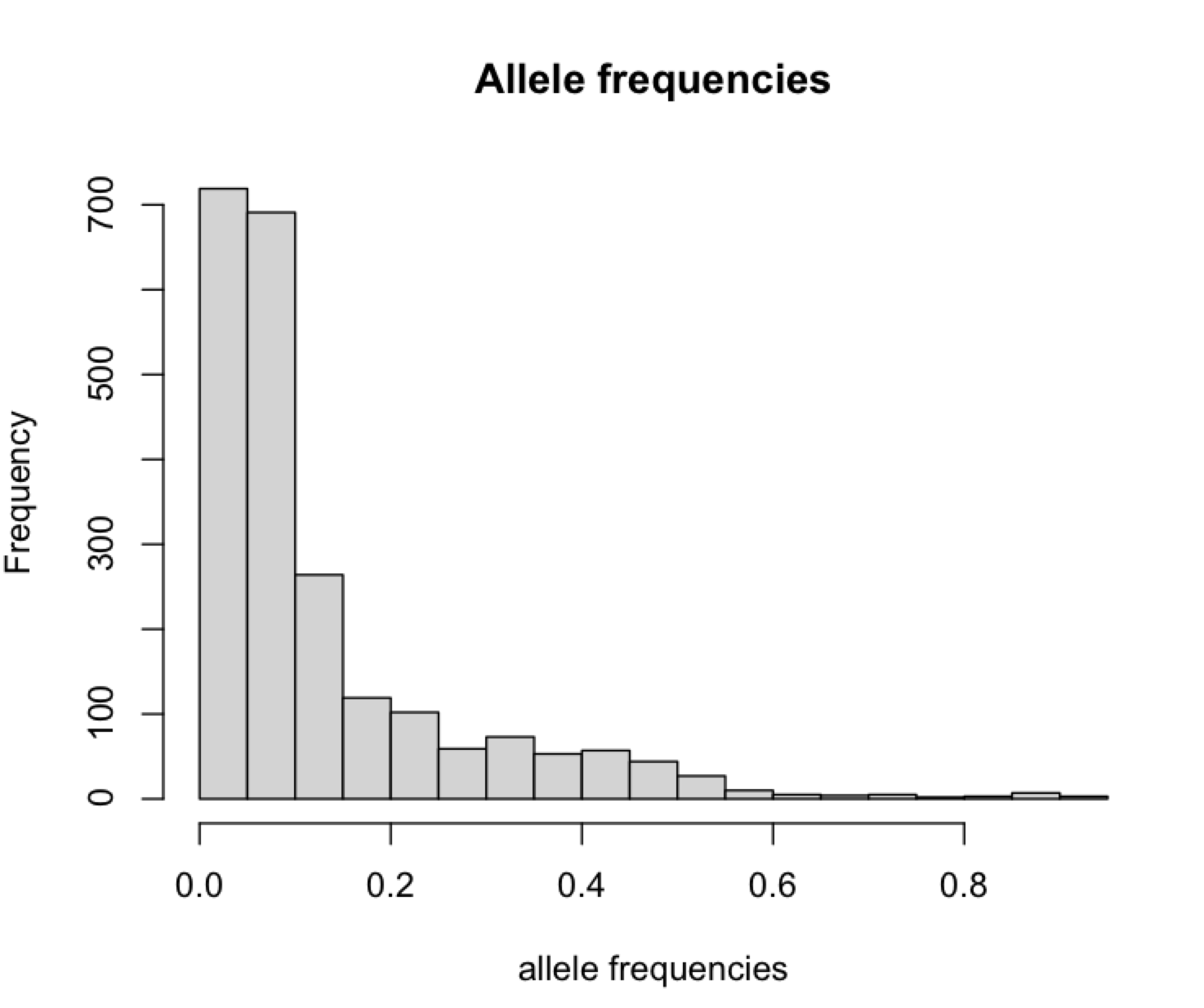

**Supplement Figure 2:** Allele frequency spectrum of remapping Illumina data to our newly established assembly GCA_019844115.1 showing high agreement and little variability across low frequency variants.

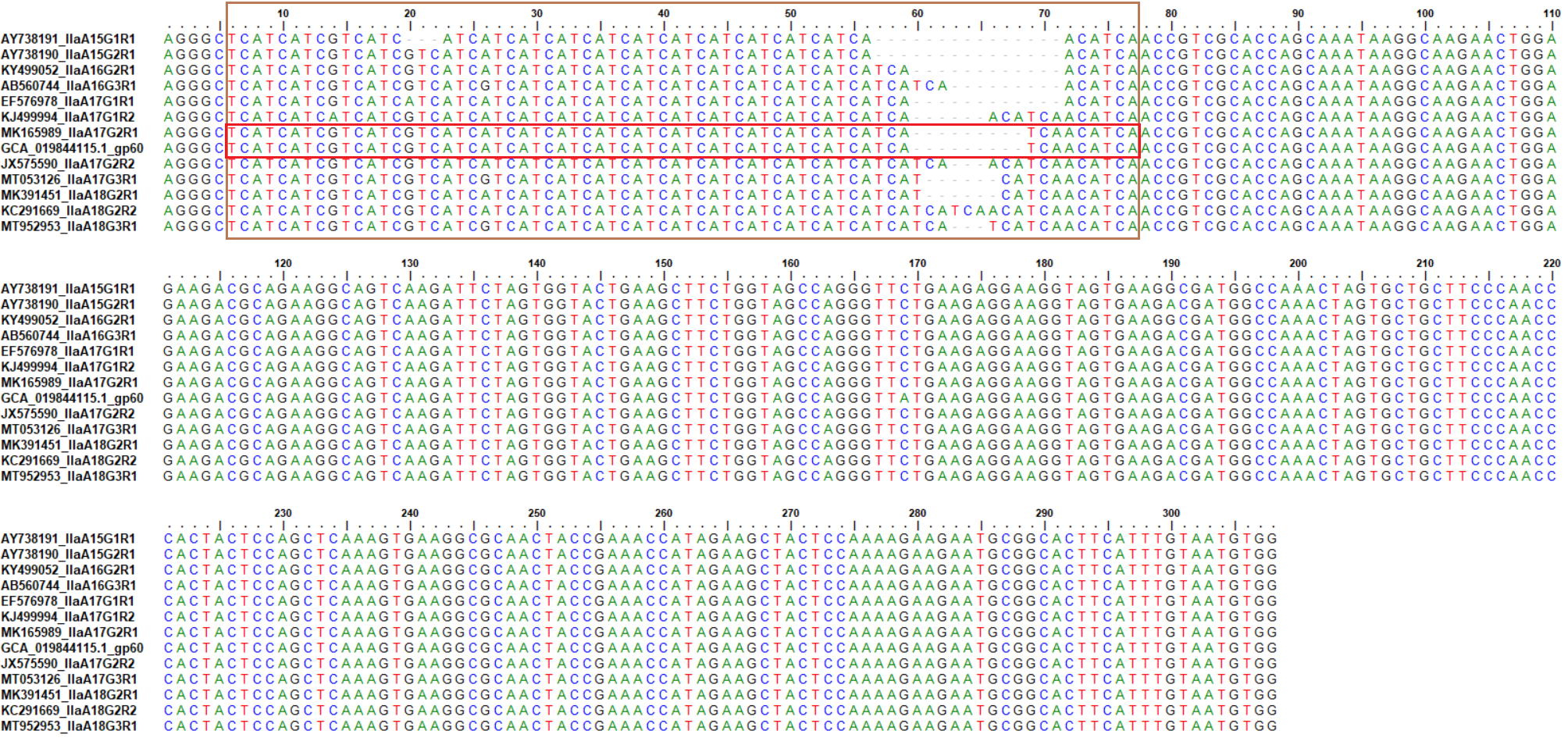

**Supplementary Figure 3**: Alignment of *gp60* subtyping region (indicated with a brown box) with the assembled genome. The genome assembly from the study (GCA_019844115.1_gp60) aligns with the reference sequence of IIaA17G2R1 (indicated with a red box, within the subtyping region).
